# Continental-scale genomic analysis suggests shared post-admixture adaptation in Americas

**DOI:** 10.1101/2020.12.16.423075

**Authors:** Linda Ongaro, Mayukh Mondal, Rodrigo Flores, Davide Marnetto, Ludovica Molinaro, Marta E. Alarcón-Riquelme, Andrés Moreno-Estrada, Nedio Mabunda, Mario Ventura, Kristiina Tambets, Garrett Hellenthal, Cristian Capelli, Toomas Kivisild, Mait Metspalu, Luca Pagani, Francesco Montinaro

**Affiliations:** Institute of Molecular and Cell Biology, University of Tartu, Tartu, Riia 23, 51010, Estonia; Estonian Biocentre, Institute of Genomics, Tartu, Riia 23b, 51010, Estonia; Department of Medical Genomics, GENYO. Centro Pfizer - Universidad de Granada - Junta de Andalucía de Genómica e Investigación Oncológica, Av de la Ilustración 114, Parque Tecnológico de la Salud (PTS), 18016, Granada, Spain; National Laboratory of Genomics for biodiversity (LANGEBIO), CINVESTAV, Irapuato, Guanajuato 36821, Mexico; Instituto Nacional de Saúde, Distrito de Marracuene, Estrada Nacional N°1, Província de Maputo, Maputo, 1120, Mozambique; Department of Biology-Genetics, University of Bari, Bari, 70126, Italy; Department of Genetics, Evolution and Environment and UCL Genetics Institute, University College London, London WC1E 6BT, UK; Department of Zoology, University of Oxford, Oxford, UK; Department of Chemistry, Life Sciences and Environmental Sustainability, University of Parma, Parma, Italy; Department of Human Genetics, KU Leuven, Herestraat 49 - box 602, B- 3000, Leuven, Belgium; Department of Biology, University of Padua, Padua, Italy

**Author notes:** These authors contributed equally as Senior authors (Luca Pagani, Francesco Montinaro).

## Abstract

American populations are one of the most interesting examples of recently admixed groups, where ancestral components from three major continental human groups (Africans, Eurasians and Native Americans) have admixed within the last 15 generations. Recently, several genetic surveys focusing on thousands of individuals shed light on the geography, chronology and relevance of these events. However, despite the fact that gene-flow could drive adaptive evolution, it is not clear whether and how natural selection acted on the resulting genetic variation in the Americas.

In this study, we analysed the patterns of local ancestry of genomic fragments in genome-wide data for ∼6,000 admixed individuals from ten American countries. In doing so, we identified regions characterized by a Divergent Ancestry Profile (DAP), in which a significant over or under ancestral representation is evident.

Our results highlighted a series of genomic regions with Divergent Ancestry Profiles (DAP) associated with immune system response and relevant medical traits, with the longest DAP region encompassing the Human Leukocyte Antigen locus. Furthermore, we found that DAP regions are enriched in genes linked to cancer-related traits and autoimmune diseases. Then, analyzing the biological impact of these regions, we showed that natural selection could have acted preferentially towards variants located in coding and non-coding transcripts, and characterized by a high deleteriousness score.

Taken together, our analyses suggest that shared patterns of post admixture adaptation occurred at continental scale in the Americas, affecting more often functional and impactful genomic variants.

## Background

The genomic variation of a substantial proportion of the individuals living in the Americas is the result of admixture involving Native American, European and African populations, together with minor recent contributions from Asia, as the results of deportation and mass migrations followed by admixture episodes [1–4].

Although many studies uncovered the complexity of the admixture dynamics in the continents [1,3–5], addressing the role of adaptive introgression in shaping the modern-day variation of American populations has been particularly challenging. In fact, the high variance in individual continental ancestries, due to very recent and still ongoing admixture, makes it hard to apply classical natural selection tools based on the distribution of genetic variation along the genome [6–9]. Following the advent of high-throughput sequencing and development of statistical tools aimed at inferring the ancestry of specific genomic regions, a commonly used approach to tackle this question focuses on loci showing ancestral proportions across the entire population that significantly diverge from whole-genome estimates [10]. Regions enriched or depleted for a given ancestry are usually interpreted as targets of natural selection, driven by post admixture adaptive pressures. Despite the large number of studies harnessing these methods, the results have not been consistent and were often not replicated. For example, the analysis of ∼2000 African Americans found putatively enriched loci (deviating more than three standard deviations) for European and African ancestries [11]. However, a subsequent study analyzing ∼29,000 individuals and applying a genome-wide significance threshold did not find any statistically significant diverging loci, highlighting the possibility of false-positive detection when less conservative thresholds are applied [12].

Furthermore, most of the selection scans performed so far focused either on single populations or on multiple groups of small sample sizes, increasing the chance of collecting false positives.

Nevertheless, some investigations of local ancestry tracts distributions in individuals from Peru, Puerto Rico, Mexico, and Colombia are concordant in suggesting rapid natural selection in genomic loci associated with immune response, such as the Human Leukocyte Antigens (HLA) [13–18].

In this study, we analyze local ancestry tracts distribution for 5,828 American individuals from 19 populations to elucidate the role of post-admixture selection in shaping the genetic variation of the Americas.

In doing so, we focused only on signals shared across multiple populations, reducing the inference of false-positives and at the same time highlighting shared or convergent episodes of selection as proposed in Yelmen et al. 2019 [19]. Moreover, given the Columbian Exchange phenomenon, the asymmetric resistance to pathogens such as Measles and Smallpox in Europeans and not in Americans, and Syphilis in Americans but not in Europeans, we specifically looked for signals in genes associated with the biology of these diseases [20–22].

Our results highlighted a series of genomic regions with Divergent Ancestry Profiles (DAP) associated with relevant medical traits such as the HLA, and others with recurring association to cholesterol and triglyceride levels, systemic lupus erythematosus and blood protein levels. Furthermore, we found that the SNPs belonging to DAP windows are enriched in genes linked to cancer-related traits and autoimmune diseases. Lastly, the analysis of the functional impact and annotation of the DAP windows revealed a larger role of natural selection on variants located in coding and non-coding transcripts and characterized by a stronger annotated impact.

## Results

### Evaluation of signals identified in multiple populations

Our aim was to identify genomic regions showing significant deviations in local ancestry assignment from the average ancestry proportion in a given population. We applied the Local Ancestry RFMix software [23] on data from the Americas as presented in Ongaro et al. (2019) [1], in which, the ancestry composition of the admixed individuals was deconvoluted using four putative source groups (Africa: 190 individuals; Europe: 289 individuals; Americas: 67 individuals; Asia: 213 individuals; Supplementary Table 1A). For simplicity we have here excluded the latter, given the low proportion of Asian ancestry detected in our previous analysis [1].

We evaluated the local ancestry output using two differently assembled datasets, as described in the Methods section. From this point on we performed all the analyses for each ancestry separately. Briefly, for the first one (20Pop dataset), we focused only on populations with more than 20% of a given ancestry; in the second one (1090Ind dataset) we excluded individuals characterized by extreme proportions for any given ancestry (< 10% and > 90%).

For the populations included in the two datasets, we estimated the ancestry specific Z-score for each SNP (Supplementary Table 2 and 3), as explained in the Methods section.

We aggregated the SNP-based output in 100kb non-overlapping windows, retaining only windows containing more than five SNPs, and annotated as DAP (Divergent Ancestry Profile) those with at least one SNP with a significant Z score (>|3|). In order to reduce the number of false positives, and focusing on shared or convergent signatures, we selected only DAP signals replicated in at least two populations. Consecutive 100kb DAP windows were grouped, and are hereafter referred to as DAP regions.

The populations and the number of haplotypes retained in the two datasets are reported in Supplementary Table 1B.

### African DAP

We identified only a single genomic window with an African divergent ancestry profile present in more than one population (Figure 1 and Table 1). This DAP window (chr15: 65,613,654-65,695,283) contains 10 SNPs within two immunoglobulin genes: *IGDCC3* and *IGDCC4*. These SNPs show significant underrepresentation of African ancestry in ACB (Barbados Island) and Dominicans when analyzed in the 1090Ind dataset.

**Figure 1:**
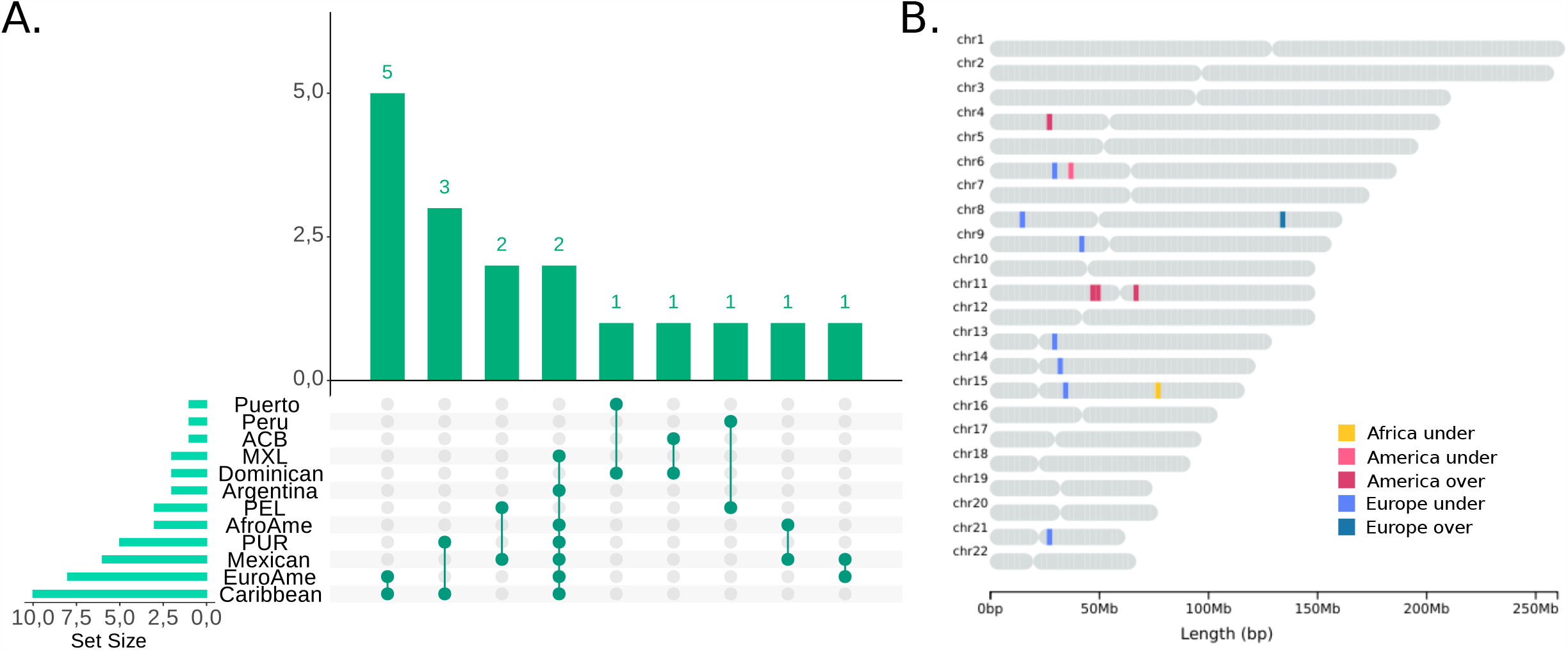
An overview of Divergent Ancestry Profile inferred by local ancestry profiles for European ancestry. A) Population distribution of European DAPs. The x axis shows DAPS in single populations while y axis shows DAP sharing among groups. This plot was built using the UpsetR package in R [48]. B) Genomic location of shared DAPs. Different colours refer to the ancestry and direction of the divergence, as indicated in the legend. The chromosome map was obtained using the chromoMap package in R.

**Table 1.**
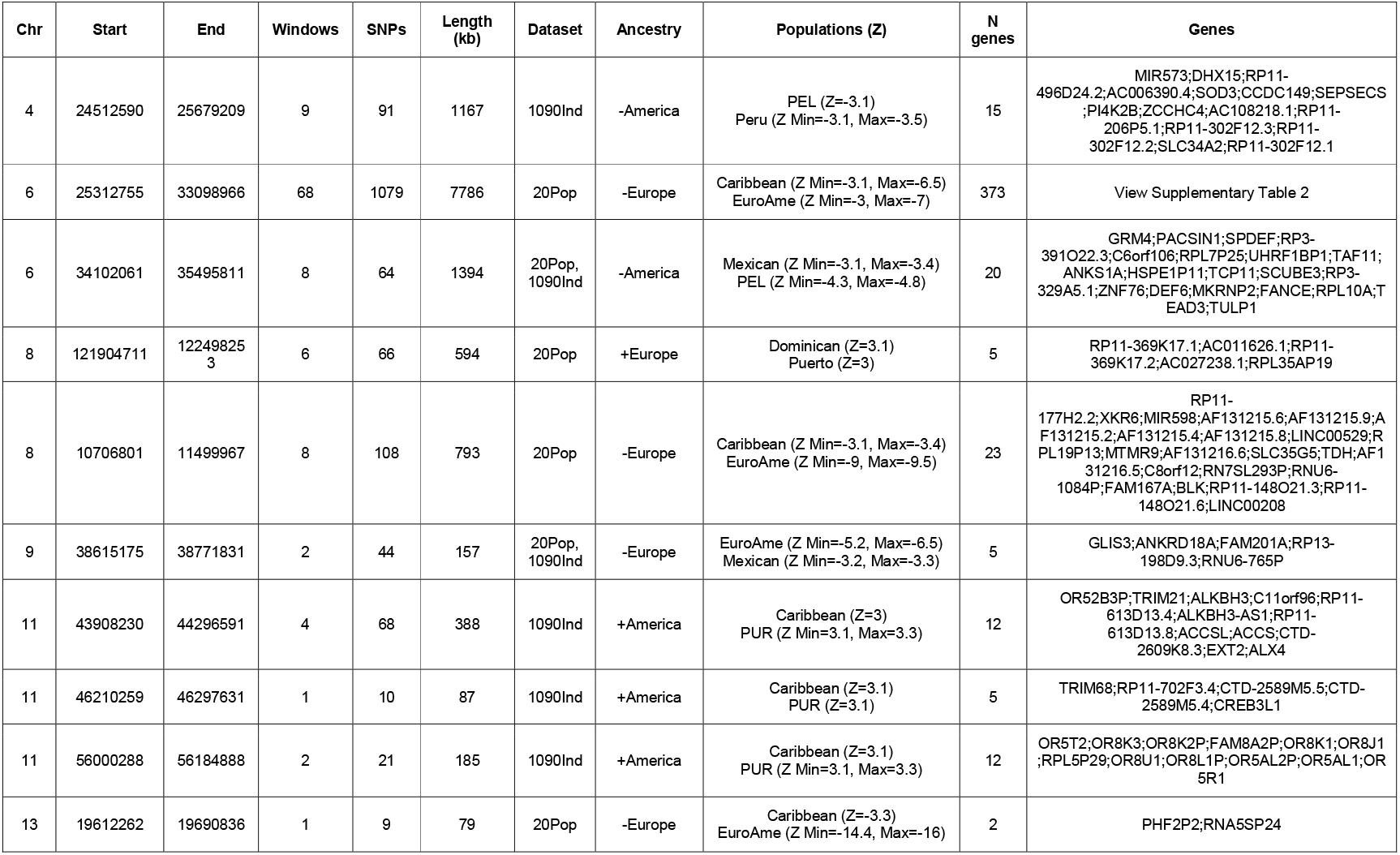
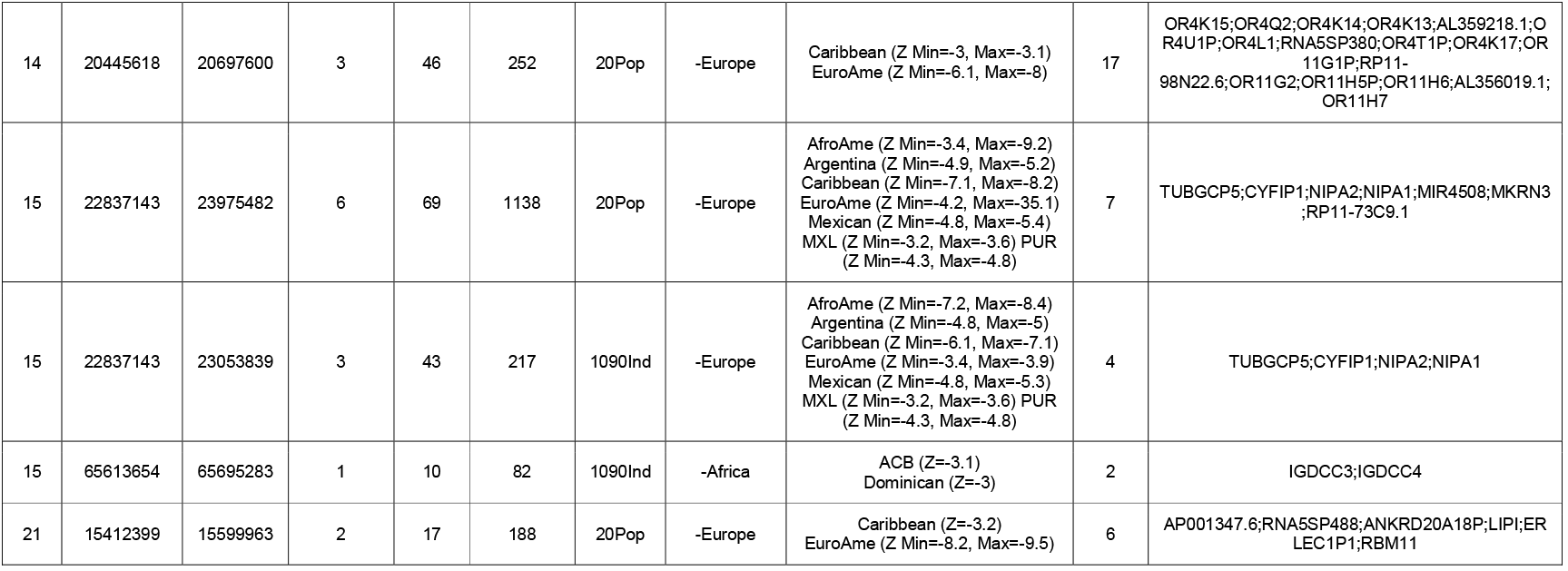
Details about inferred genomic regions with Divergent Ancestry Profiles. “Windows” refers to the number of DAP 100kb windows inside the genomic region; “SNPs” shows the number of SNPs contained in the genomic region; “Ancestry” refers to the specific ancestry for which we found an underrepresented (-) or overrepresented (+) DAP window or region; “Populations (Z)” reports Z scores for the significant populations; “N genes” shows the number of genes related to SNPs part of the genomic region; “Genes” refers to the Gene/clone identifiers provided in ENSEMBL annotation.

### European DAPs

We identified 7 DAP genomic regions with underrepresented European ancestry in the 20Pop dataset, of which 2 are replicated in the 1090Ind dataset; in addition, we found one overrepresented region in the 20Pop dataset. More details about the content of the DAP genomic regions are reported in Table 1.

Strikingly, the major signal was found for a region in chromosome 6 extending for a total of 7.8 Mb (chr6:25,312,755-33,098,966) with underrepresented European ancestry in two populations, Caribbean and European Americans (EuroAme). This region contains 373 genes (Supplementary Table 4) including the large locus of the Human Leukocyte Antigen (HLA). Nineteen of the 1079 SNPs typed in our dataset are predicted to have a high functional impact, according to the Combined Annotation Dependent Depletion (CADD) annotation (PHRED-scaled C-score > 20, Supplementary Table 5, PHRED hereafter). Interestingly, six of the 100kb windows included in this region carry high proportions of SNPs with large PHRED values (>= 20; 26% of all SNPs in chr6:31,602,967-31,688,217, 10% in both chr6:26,413,088-26,484,376 and chr6:29,407,970-29,483,911, 8.3% in both chr6:28,300,336-28,391,465 and chr6:32,019,769-32,083,175 and 7.7%in chr6:30,823,630-30,897,022), corresponding to the top 5% of the distribution estimated on all the genomic windows (26% corresponds to 0.0006, 10% to 0.025, 8.3% to 0.039 and 7.7% to 0.039). Then, the variants in this DAP region belong to 35 different genes, of which 32 are related to HLA.

We found several SNPs in this region with significant disease associations in the GWAS catalog. For example, the SNP with the highest value of PHRED (PHRED=46, associated with the top ∼10^−4^ quantile of the genome-wide distribution, Supplementary Table 5), *rs2071543*, (chr6:32,811,629) is associated with IgA glomerulonephritis (also called Berger Disease); it has been reported as genome-wide significant in East Asians (p-value = 2.1 × 10^−9^), but not in Europeans (p-value = 0.85) [24]. In our dataset, the mean frequency of the risk allele (G) in the Native American sources is 0.76, while in Caribbean and European Americans is 0.89 and 0.88, respectively.

Then, we found the *rs8111* (*ATF6B* gene, PHRED=22.2, Supplementary Table 5) associated with Blood Protein Levels in European populations [25]; this association has been found also for other seventeen SNPs reported in the GWAS catalog [26–29].

In addition, a variant (*rs3130618, PHRED=32)* belonging to *GPANK1* gene is associated with MMR (measles, mumps and rubella) vaccination-related febrile seizure [30] and Membranous Glomerulonephritis [31]. Inside this region, two SNPs, *rs140973961* and *rs78331658*, not present in our genome-wide dataset, have been associated with measles and immune response to the measles vaccine, respectively, in the GWAS catalog [32,33]. To the best of our knowledge these are the only results that might be related to the phenomenon of Columbian exchange.

Notably, we observed a neighboring region (chr6:34,102,061-35,495,811), distant ∼1Mb from the one reported above, with underrepresented American ancestry for 20Pop and 1090Ind, as shown in Table 1. In both cases we found the same region spanning 1.8 Mb and containing 64 SNPs underrepresented in both Mexican and PEL. In this region, *rs2395617* (chr6: 35,285,720; PHRED=19.6, Supplementary Table 5) in the *DEF6* gene, has been associated with Apolipoprotein A1 levels in European individuals [34]. Similarly, another variant (*rs2814982*, chr6:34,546,560) has been reported associated with cholesterol levels and LDL levels in multiple studies [35–37].

We registered a similar region, 1.1 Mb (chr6:34,332,179-35,495,811), with an overrepresentation of African ancestry (20Pop dataset) in Caribbean (Min Z= 3.2, Max Z= 3.4) and with values close to the significance threshold in the African Americans (AfroAme, Min Z=2.4, Max Z=2.6).

We observed, in both the analysed datasets (although characterized by different size), a signal of underrepresented European ancestry in chromosome 15 (Figure 1 and Table 1). Among the others, a SNP (*rs2278458*, chr15:22,999,857) in *CYFIP1* gene is associated with Triglyceride levels and to the response to diuretics in individuals of European and African ancestry [38]. Strikingly, this DAP region is shared across seven populations: African Americans (AfroAme), Argentina, Caribbean, European Americans (EuroAme), Mexican, MXL and PUR.

The remaining four genomic regions with significantly lower European ancestry in Caribbean and European American individuals were observed only in 20Pop dataset and are located in chromosomes: 8, 13, 14 and 21 (Supplementary Text).

Additionally, we identified a DAP region in European ancestry of the 20Pop dataset that is overrepresented in Dominican and Puerto Ricans (Puerto, Table 1). This region is in chromosome 8 and contains the *rs4871180* (chr8:122,259,074) that could be associated with Diverticular disease in British individuals [39]. The frequency of the risk allele (T) of this variant is very high in the analysed Native American source populations, ranging from 0.87 to 0.9, while it has frequencies of 0.33 and 0.23 in Dominican and Puerto, respectively.

### American DAPs

When the American ancestry profiles were evaluated besides the DAP region of chromosome 6 discussed above, we identified one depleted and three enriched genomic regions in the 1090Ind dataset. Three different DAP regions with observed overrepresentation in Caribbean and Puerto Rico (PUR) were found on chromosome 11 (Figure 1, Supplementary material and Table 1). The distance between the first two regions is approximately 2Mb, and could in fact be part of the same selection event. One of these two regions (chr11:46,210,259-46,297,631; 87kb) host a *SNP* (the *rs10437653*) that has been shown to be associated with birth weight in Europeans [40].

The last region map on chromosome 4 (chr4:24,512,590-25,679,209, Table 1 and Figure 1) and is underrepresented for American ancestry in two Peruvian populations, PEL and Peru. Interestingly, we identified two SNPs within this region that are associated with diseases/traits: *rs12500612* (chr4:24,740,958) with Major depressive disorder in Europeans [41]; and *rs1395221* (chr4:24,626,903), with Apolipoprotein A1 and HDL cholesterol levels in European individuals [34]. The mean frequency of the risk allele (G) of *rs1395221* in the analysed Native American source populations is 0.90 and 0.60 in the European ones; whereas PEL and Peru have a frequency of 0.77 and 0.78, respectively.

### The functional impact of DAP regions in the Americas

We explored the functional impact measured as SNP deleteriousness (PHRED-scaled C-score) and functional annotation (Annotype), both derived from the CADD annotation, focusing on the multiple population signals described above.

When we compared the distribution of all the PHRED values belonging to windows showing divergent ancestry profiles (DAP) with the non-divergent ones (non-DAP), for European ancestry values were significantly higher in SNPs from DAP windows than those from non-DAP (Wilcoxon test, p-value=2.8E-29). The same result was observed for the American ancestry (Wilcoxon test, p-value=0.00014) of the 20Pop dataset (Figure 2 and Supplementary Table 6A). No comparison within the 1090Ind dataset showed any significance (Supplementary Figure 1).

**Figure 2:**
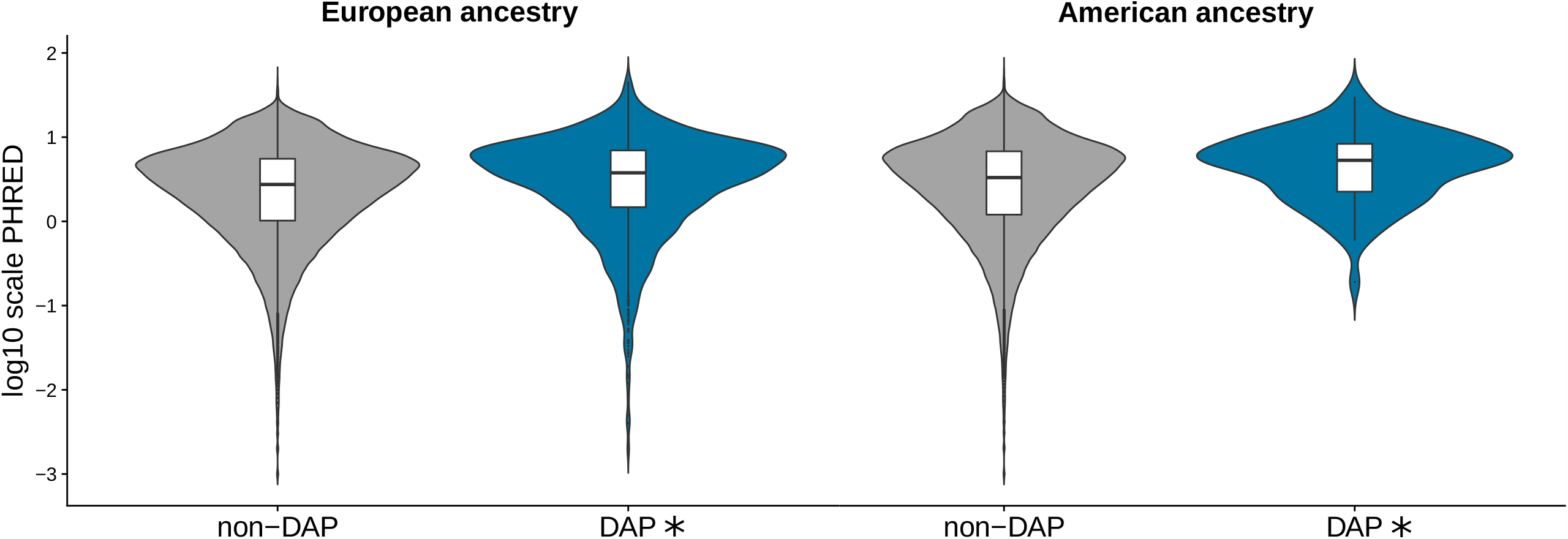
Comparison of the distribution of all the PHRED-scaled C-score values belonging to windows with divergent ancestry profiles (DAP) with the ones from the non-divergent for European (96 DAP and 16,760 non-DAP windows) and American ancestries in the 20Pop (8 DAP and 16,848 non-DAP windows) dataset. The asterisk refers to a statistically significant p-value (Wilcoxon test, alpha=0.01). The number of analysed windows is reported in Table 6A.

Then, in DAP and non-DAP windows, we analysed the distribution of the proportion of the five types of functional annotation: Coding Transcript (variants in protein-coding exons), Untranslated Transcript (variants in UTRs and introns), Non-Coding Transcript (mature miRNA and non-coding transcript exon variants), Regulatory Feature and Intergenic. In detail, for each 100kb window, we annotated the relative functional composition of all the SNPs, and evaluated the proportion of each category (Figure 3, Supplementary Figure 2 and Supplementary Table 6B). Every SNP was assigned to at least one Annotype.

**Figure 3:**
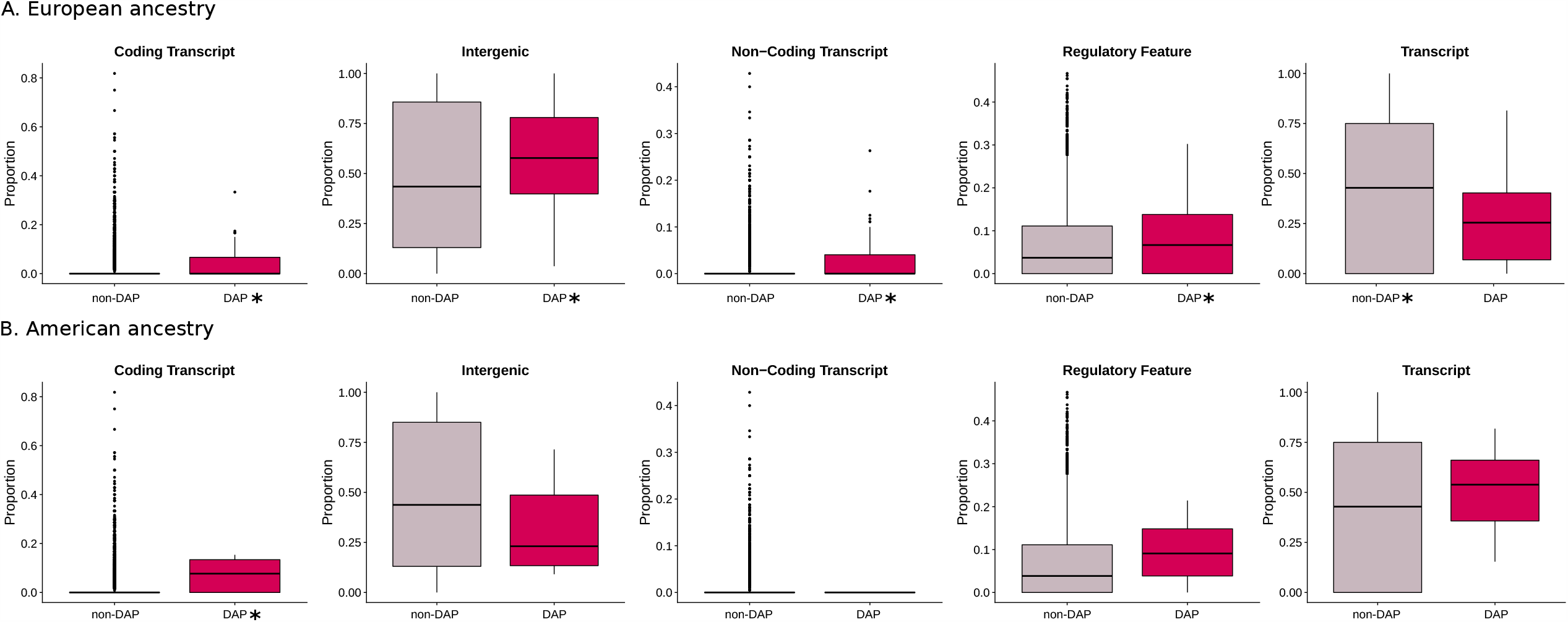
Comparison of the distribution of Annotypes (Coding Transcript, Intergenic, Non-coding Transcript, Regulatory Feature, Transcript) belonging to divergent ancestry profiles (DAP) windows with the ones from the non-divergent for European (A, 96 DAP and 16,760 non-DAP windows) and American (B, 8 DAP and 16,848 non-DAP windows) ancestries in the 20Pop dataset. The asterisk refers to a statistically significant p-value (Wilcoxon test, alpha=0.01). The number of analysed windows is reported in Table 6B.

In detail, for each 100kb window, we annotated the relative functional composition of all the SNPs, and evaluated the proportion of each category (Figure 3, Supplementary Figure 2 and Supplementary Table 6B). Each SNP was assigned to at least one Annotype.

We observed that the proportion of “Coding transcript”, “Non-coding Transcript”, “Intergenic” and “Regulatory Feature” is higher in DAP windows than non-DAP in the European ancestry (20Pop Dataset, Wilcoxon test, p-values reported in Supplementary Table 6B, Figure 3). In contrast, the proportion of “Untranslated Transcript” is higher in non-DAP SNPs than DAP (European ancestry 20Pop Dataset, Wilcoxon test, p-value=0.00014, Figure 3).

Moreover, when we looked at the Native American ancestry the proportion of “Coding Transcript” was higher in DAP SNPs than in non-DAP for both datasets (Wilcoxon test, p-value=0.0025, Figure 3).

### Gene Set Enrichment Analysis

We performed gene set enrichment analysis (GSEA) for the genes encompassing the DAP SNPs, in order to understand if specific health/disease pathways or phenotypes are significantly enriched or depleted in the continental ancestries of multiple populations. We explored five different libraries: Human 2019 Kyoto Encyclopedia of Genes and Genomes (KEGG), Gene Ontology (GO) 2018, GTEx Tissue Sample Gene Expression Profiles down and up and Genome-wide Association Studies (GWAS) Catalog 2019.

Interestingly, we found signals exclusively related to underrepresented variants in the European ancestry in the 20Pop dataset from four libraries (GWAS, KEGG, GO and GTEx up) and related to overrepresented variants in the Native American ancestry in the 1090Ind dataset from only one library (KEGG).

In detail, for the European ancestry results, we found 46 significantly overrepresented traits in GWAS Catalog, where the most significant one is related to the *Autism spectrum disorder or schizophrenia* (p-value=3.9E-217) term followed by *Blood protein levels* (p-value=4E-84) and *Ulcerative colitis* (p-value=7.7E-49). Six significant terms are associated with types of cancer: *Lung cancer* (p-value=1.9E-33), *Squamous cell lung carcinoma* (p-value=2.7E-31), *Lung cancer in ever smokers* (p-value=1.3E-29), *Small cell lung carcinoma* (p-value=4.1E-08), *Prostate cancer* (p-value=0.00011) and *Cervical cancer* (p-value=0.00016). Four are connected with Hepatitis B, both related to the chronic infection and to the response to the vaccine. Then, a group of significant traits is linked to autoimmune diseases such as *Sarcoidosis, Psoriasis, Lupus, Behcet’s disease, Type I diabetes and Inflammatory Bowel disease*.

Eighteen traits are overrepresented when the KEGG library is explored, where the *Systemic lupus erythematosus* (p-value=2.6E-46), that is in common with GWAS library list of terms, has the lowest adjusted p-value followed by *Alcoholism* (p-value=7.8E-27) and *Viral carcinogenesis* (p-value=6E-14). Other terms in common with the GWAS catalogue are *Inflammatory Bowel disease* (KEGG: p-value=0.00022, GWAS: p-value=2.7E-29) *and Type I diabetes* (KEGG: p-value=1.4E-11, GWAS: p-value=5.9E-19).

Finally, significant GO enrichment was found for *MHC class II receptor activity (*p-value=2.29E-05) and for GTex *lung expression in females of 60-69 years, (*p-value=0.00036). The results are reported in Figure 4, where only the first ten most significant terms for each annotation library are shown, while in Supplementary Table 7A-D all the results are reported.

**Figure 4:**
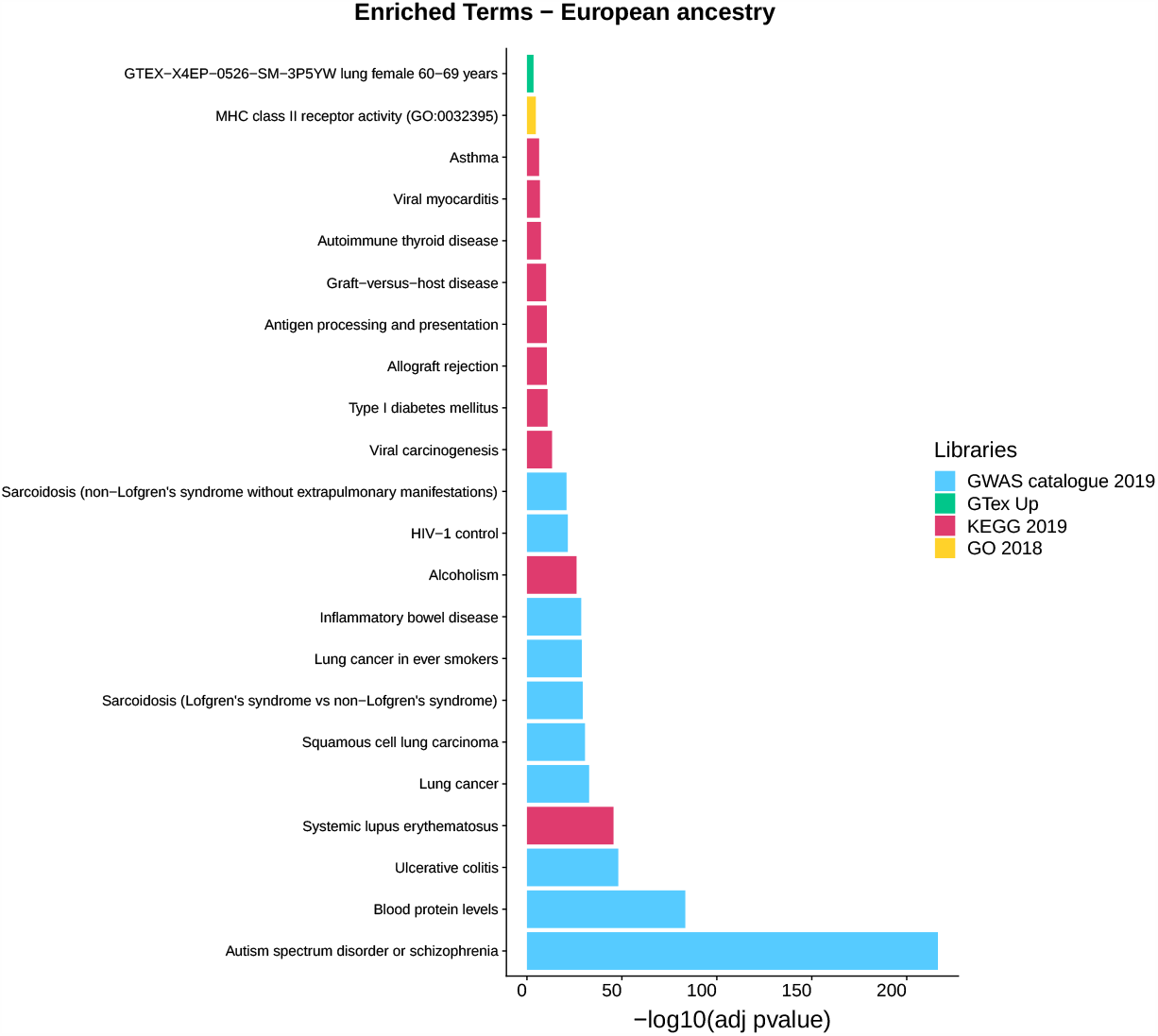
Gene-set enrichment analysis results for the European ancestry of the 20Pop dataset (Supplementary Table 7A-D). Only the first ten enriched terms for library are shown. Libraries: Genome-wide Association Studies (GWAS) Catalog 2019, GTEx Tissue Sample Gene Expression Profiles up, Human 2019 Kyoto Encyclopedia of Genes and Genomes (KEGG) and Gene Ontology (GO) 2018.

When we excluded the gene list related to the chromosome 6 DAP region with underrepresented European ancestry the GSEA gave no significant results. For the Native American ancestry, we identified one significant enriched term related to *Olfactory Transduction* (p-value=0.00047) according to the KEGG library (Supplementary Table 7E).

## Discussion

In this study, we evaluated the presence of post-admixture regions with a divergent ancestry proportion on a broad dataset composed by 5,828 individuals from 19 recently admixed populations. As previous studies have focused on small sample sizes and a single population, our approach aimed at the identification of repeated signals at a Pan-American scale, allowing at the same time, the reduction of false positive rate, and the discovery of selective forces acting at the scale of the entire continent. Furthermore, we took into consideration possible errors by the LAI algorithm harnessed here, performing the same analysis discarding individuals with extreme values of a given ancestry.

In doing so, despite the potential for African admixtures to enrich the genetic diversity of any recipient human group, among the five populations in the 20Pop dataset and the seven populations in the 1090Ind dataset having a relatively high contribution from Africa, we identified only a single underrepresented and no overrepresented region. Although this scenario is in contrast with surveys identifying DAP regions of African inheritance [17,18], our results might be explained by the high degree of conservativeness of our approach. Moreover, the recently documented high genetic variation of the African populations contributing to the modern-day American populations, coupled with the relatively low African ancestry proportion in the analysed samples, may have provided differential bases for the subsequent selective forces [1,42–44].

On the other hand, we identified 7 underrepresented regions which consistently showed deviation from the ancestral proportions of the European ancestry, including a ∼8Mb region in chromosome 6 encompassing the Major Histocompatibility Complex II. A neighboring region was identified as underrepresented also for the American ancestry, and although we failed to identify a replicated signal for African ancestry, it is worth noting that the HLA region identified as underrepresented for the European ancestry has been found overrepresented for the African ancestry in Caribbean, and close to the significant threshold in African American, confirming previous results [13–18].

Replicated signals for overrepresented regions that were found in the Americas are consistent with a scenario in which the 15,000 years of the peopling of the continents resulted in adaptation.

Our evaluation of the biological impact of post-admixture adaptation in the Americas revealed that, irrespective of their direction, SNPs having high functional or phenotypic consequences tend to be “selected” more often than those with mild effect. This is also confirmed by the fact that SNPs in UTRs and introns are not a preferential target of natural selection, in contrast to what has been observed for SNPs in coding and regulatory regions (Figure 3 and Supplementary Figure 2).

Lastly, the gene set enrichment analysis performed here revealed that selection acted predominantly on regions associated, among the others, to the onset of autoimmune disease, various protein levels in blood and several different kinds of cancer. The fact that we did not obtain significant results when we removed from the GSEA the underrepresented DAP region in chromosome 6 of European ancestry could be a sign of reduced polygenic enrichment in our data.

### Conclusions

Overall, our research suggests that common selective pressure in the Americas had a non-negligible impact on shaping the genetic variation of the two continents, while the Columbian Exchange phenomenon seems to have played just a minor role. Our results also indicate that given the limitations of genetic scan for natural selection algorithms implemented so far, the analysis of multiple population datasets characterized by high sample size will be essential for both the identification and characterization of post admixture adaptation at a more local scale.

## Material and Methods

### Genome-wide data

The analysed genome-wide dataset was recovered from Ongaro et al 2019 [1]. This dataset was filtered using PLINK ver. 1.9 [45] to include only SNPs and individuals with genotyping success rate > 97%, retaining a total of 251,548 autosomal markers. For this study, we used the reference genome version b37.

We removed 22,295 SNPs belonging to centromeric regions or to the first or last 5 Mb of the chromosomes on the basis of the information retrieved from UCSC browser (https://genome.ucsc.edu/cgi-bin/hgTables); following this step we kept a total of 229,253 variants. In the current study we analysed 6,587 individuals, of which 5,828 belong to 19 admixed American populations while the remaining 759 samples are from populations from Africa, Asia, Europe and America. The first set of individuals represents the so-called “targets”, while the second set represents the “sources”. The details are reported in Supplementary Table 1A-C.

### Phasing

Germline phase was inferred using the Segmented Haplotype Estimation and Imputation tool (ShapeIT2) software[46], using the HapMap37 human genome build 37 recombination map.

### Local ancestry

We estimated local ancestry assignation for genomic fragments of the target American individuals with RFMIX software[23], using the following reference source populations: Yoruba (YRI), Gambia (GWDwg) and Mozambique for Africa, Chinese Han (CHB) and Japanese (JPT) for Asia, Spanish (IBS), British (GBR) and Tuscany (TSI) for Europe and Tepehuano, Wichi and Karitiana for Native American ancestry (Supplementary Table 1A). We used “PopPhased”, “-n 5” and “--forward-backward” options as recommended in the RFMix manual.

Starting from the RFMix output files we built four PLINK file sets, one for each of the four source ancestries, masking in each one the SNPs assigned to any of the other three ancestries. In details, an allele was assigned as missing in the PLINK file of ancestry A when that allele was not assigned to ancestry A in the “Viterbi” output file or the probability of belonging to ancestry A (as reported in the “forward-backward” output) was less than a defined threshold (< 0.9). In this and in the following analyses we considered the samples as separated into the two phased haplotypes.

Given that Asian ancestry was consistently found at low frequency (range:0.2-9.4%) we decided to not include this ancestry in our search for DAP windows. At this point we adopted two different analysis approaches and assembled two datasets (Supplementary Table 1B):

1. 1090Ind dataset: this dataset was assembled considering the ancestral proportions of each individual. In detail, we used the PLINK command --missing to obtain a missing report for each individual; then using the information from “.imiss” files we kept those individuals with more than 0.1 (10%) and less than 0.9 (90%) of F_MISS (representing the missingness of the individual) for each ancestry
2. 20Pop dataset: this was assembled taking in consideration the ancestral proportions averaged by populations. In this case we kept for the analyses only those populations with at least 20% of a specific ancestry. These proportions were recovered from the global ancestry analysis (SOURCEFIND [4]) reported in Ongaro et al 2019 [1].

### SNPs annotation

We annotated all our SNPs with CADD (Combined Annotation Dependent Depletion, [47]), a tool for scoring the deleteriousness of single nucleotide variants in the human genome.

### Detecting deviation from the expected ancestral proportions

In order to detect deviation from the expected ancestral proportions in Local Ancestry Inference, we estimated the per-SNP average population assignation.

In doing so, starting from the ancestry-specific PLINK files described above, we initially calculated the population specific missingness using --missing --family in PLINK 1.9 [45]. We then estimated the proportion of Local ancestry assignment as 1-missing. The obtained proportions were finally standardized calculating the Z-score for each SNP. Finally, we partitioned each chromosome into windows of 100,000 bp, obtaining 16,857 windows, and defined as Divergent Ancestry Profile (DAP) windows the ones that contained more than five SNPs of which at least one presented a significant Z-score. We decided to set as statistically significant values of |Z|>3 (Supplementary Table 2-3). In order to further reduce the identification of false positives, we focused exclusively on signals replicated in at least two populations.

### PHRED-scaled C-score analysis

We explored the differences in values of PHRED in the so-called DAP and non-DAP windows. We first extracted from the 100kb windows all the SNPs with a corresponding PHRED value retrieved from the CADD annotation. Then, we compared the distribution of all the PHRED values belonging to the DAP windows with the ones from the non-DAP using a paired Wilcoxon test with R.

### Annotype analysis

We investigated the differences in the distribution between DAP and non-DAP windows of the five types of functional annotations (Annotype; Coding Transcript, Intergenic, Non Coding Transcript, Regulatory Feature and Transcript). In CADD annotation, the broad category of transcripts is divided into Coding Transcript, Non-Coding Transcript and Transcript. In details, “Coding Transcript” refers to several types of variants like missense, synonymous, stop-gained, stop-lost, initiator codon, stop-retained, frameshift, inframe insertion, inframe deletion, incomplete terminal codon and protein altering. The “Non-Coding Transcript” category contains mature miRNA and non-coding transcript exon variants, while “Unspecific Transcript” refers to UTRs and introns.

In detail, we extracted all the SNPs from the windows of interest recovering the corresponding Annotype, then we estimated the Annotype proportions for each window (DAP and non-DAP) and we compared the proportion distributions using a paired Wilcoxon test with R.

### Gene set enrichment analysis (GSEA)

GSEA was performed using the GSEAPY enrichr python module on gene lists obtained from the results of the detection of the DAP windows. We focused on the over and underrepresented SNPs from the replicated DAP windows related to the three continental ancestries; then, we extracted the genes assigned to those SNPs by the CADD annotation. In detail, we compiled 7 different gene lists to use as inputs for GSEA. These lists were extracted from multiple population signal results and assembled as follows: a) 20Pop and 1090Ind European ancestry negative (Z < −3) DAP SNPs, b) 20Pop European ancestry positive (Z>3) DAP SNPs, c) 20Pop and 1090Ind Native American ancestry negative DAP SNPs, d) 1090Ind Native American ancestry positive DAP SNPs and e) 1090Ind African ancestry negative DAP SNPs.

Only the combinations of datasets and ancestries with at least one DAP window are present. We used five libraries:

- Human 2019 Kyoto Encyclopedia of Genes and Genomes (KEGG) that is a database resource for understanding high-level functions and utilities of the biological system.
- Gene Ontology (GO) Molecular Function 2018.
- Genotype-Tissue Expression (GTEx Tissue Sample Gene Expression Profiles down and up) both for down and up regulated genes.
- Genome-wide Association Studies (GWAS) Catalog 2019.

We applied a significance cutoff threshold of 0.001, and any adjusted p value below this cutoff was therefore considered as significant.

### Supplementary tables captions

**Supplementary Table 1**. Details of the genotype data used in this study at population (A) and at individual (C) level. “S/R” column refers to whether the sample was used as “Source” or “Recipient” in the Local Ancestry pipeline. B) 20Pop and 1090Ind datasets details.

**Supplementary Table 2**. 20pop dataset Z scores by SNP in the three ancestries (A. Africa

B. Europe C. America).

**Supplementary Table 3**. 1090Ind dataset Z scores by SNP in the three ancestries (A. Africa

B. Europe C. America).

**Supplementary Table 4**. Genes belonging to the DAP region of chromosome 6 underrepresented in the European ancestry of 20Pop dataset. They refer to the Gene/clone identifiers provided in ENSEMBL annotation.

**Supplementary Table 5**. List of SNPs belonging to DAP genomic regions with a PHRED value (PHRED-scaled C-score) higher than 20. The information reported in columns J to R were retrieved directly from CADD annotation.

**Supplementary Table 6**. P-values derived from the comparison of the distribution of PHRED-scaled C-scores (A) and of the five types of functional annotation (B) between SNPs belonging to DAP and non-DAP windows.

**Supplementary Table 7**. Statistically significant gene-set enrichment analysis results for the European ancestry (A. Geno Ontology, B. GTex up, C. GWAS catalog, D. KEGG libraries) and for the American ancestry (E. KEGG library) of the 20Pop dataset. A term is statistically significant when the adjusted p-value is lower than 0.01 (highlighted in green).

## Authors’ contributions

LO, FM, and LP conceived, designed, and supervised the project. LO and FM conducted the analyses and generated the manuscript figures. MEA-R, AM-E, NM, CC, KT and MMe provided resources. LO, FM, LP and TK wrote the manuscript, with inputs from all the authors

## Acknowledgements

We thank the people working at the High Performance Computing Center of the University of Tartu for the help and support provided.

## Funding

This research was supported by the European Union through the European Regional Development Fund (Project No. 2014-2020.4.01.16-0030 to LO, MMe, FM; Project No. 2014-2020.4.01.16-0271 to RF; Project No. 2014-2020.4.01.16-0125 to RF; Project No. 2014-2020.4.01.16-0024, MOBTT53 to DM, LM, LP). This work was supported by the Estonian Research Council grant PUT (PRG243) (to RF, MMe, LP). This research was supported by the European Union through the European Regional Development Fund (project no. 2014-2020.4.01.16-0024 to TK). This research was supported by the European Union through Horizon 2020 grant no. 810645 (to MMe). This research was supported by the European Union through the Horizon 2020 research and innovation programme under grant no 810645 and through the European Regional Development Fund project no. MOBEC008 to MMo.

## Competing interests

The authors declare that they have no competing interests

## Notes

### Competing Interest Statement

The authors have declared no competing interest.

