## Supplemental text and figures for "Continental-scale genomic analysis suggests shared post-admixture adaptation in Americas"

**Supplementary text**

**Evaluation of signals identified in multiple populations**

**European DAPs**

The details about the content of the genomic regions with divergent ancestry profile are reported in Table 1.

More detail about the interesting SNPs reported in the main text and also other variants are described below.

In the DAP region of chromosome 6 with underrepresented European ancestry we found:

- The *rs3130453* (chr6:31,124,849, PHRED=40) is part of *CCHCR1* and *TCF19* protein coding genes, in proximity to HLA (Human Leukocyte Antigen) and it is associated with Psoriasis [[1]](https://paperpile.com/c/lkE0et/Yj0ad). Furthermore, *rs130072* (chr6:31,112,484)*,* located in the same region, is part of the *PSORS1C1* gene which has been associated with *Psoriasis* and *Systemic sclerosis [[2,3]](https://paperpile.com/c/lkE0et/RgM2Z+VBjLw)*.
- *The rs2071543* (chr6:32,811,629) is part of four different genes: PSMB8, PSMB9, *TAP1* and *TAPSAR1*. The first two, located in the class II region of the MHC, are immunoproteasome and are associated with diseases like *Proteasome-Associated Autoinflammatory Syndrome 1* and *Nasopharyngeal Disease[[4,5]](https://paperpile.com/c/lkE0et/jaFOA+x3Zxc)*, respectively. On the other hand, diseases associated with *TAP1*, a protein coding gene, include *Immunodeficiency By Defective Expression Of MHC Class I* and *Bare Lymphocyte Syndrome, Type I[[6,7]](https://paperpile.com/c/lkE0et/LhLz1+Ydazy)*. Finally, TAPSAR1 is an RNA gene affiliated with Long non-coding RNA class.
- The *rs1046089* (PHRED=25.9, Supplementary Table 5) belongs to the *PRCC2A* gene has been reported to be associated in European individuals with *Menopause (age at onset)[[8,9]](https://paperpile.com/c/lkE0et/2FcoR+zTq2S)* and *Schizophrenia[[10]](https://paperpile.com/c/lkE0et/QDaiF)*.

We identified six additional DAP regions with underrepresented European ancestry (Table 1). Two of them, in chromosome 9 and 15, were observed for both the 20Pop and 1090Ind approaches. The former is identical in both datasets, extending for 157kb and containing 44 SNPs (chr9:38,615,175-38,771,831) located in five genes. In the 20Pop dataset this DAP region is shared between European Americans (Min Z=-5.2, Max Z=-6.5) and Mexicans (Min Z=-3.2, Max Z=-3.3), while in the 1090Ind is shared between African Americans (AfroAme, Min Z=-3.2, Max Z=-3.4) and Mexicans (Min Z=-3, Max Z=-3.3), reflecting the different composition of the two datasets. Notably, *rs7039377* (chr9:38,675,465) has been associated with obesity-related traits in Hispanic children [[11]](https://paperpile.com/c/lkE0et/yTea1).

The remaining four genomic regions for which the European proportion is significantly lower have been observed only in 20Pop dataset and involve four chromosomes: 8, 13, 14 and 21; we found them underrepresented across Caribbean and European American individuals. The one in chromosome 8 (chr8:10,706,801-11,499,967) spans 793kb and contains 108 SNPs that are part of 23 genes. Of these 108 SNPs, 15 are associated with at least one trait in the GWAS catalog, of which three of them are associated with Systemic lupus erythematosus in eight different studies[[12–19]](https://paperpile.com/c/lkE0et/Bk1BN+WtqGb+FopPv+ZWal1+JX6OL+q6JUt+AoEfh+cKVsn). A DAP window of 79kb in chromosome 13 (chr13:19,612,262-19,690,836) is characterised by 9 variants belonging to two genes, PHF2P2 and RNA5SP24.

Interestingly, inside the region of chromosome 14 (chr14:20,445,618-20,697,60; 46 SNPs and 17 genes) we pointed out the *rs2775254* (chr14:20,528,528) that has a PHRED of 23.3 and is included in the OR4L1 gene (Supplementary Table 5). Moreover, an association between the metabolite levels and the *rs1188568* (chr14:20,656,645) has been reported [[20]](https://paperpile.com/c/lkE0et/BQP0T). Finally, in chromosome 21 we found a region of 188kb (chr21:15,412,399-15,599,963) with seventeen SNPs that are part of six genes (Table 1) of which LIPI is associated with Heart rate in heart failure in European and African Americans [[21]](https://paperpile.com/c/lkE0et/9bBJQ) and Neuritic plaques measurement [[22]](https://paperpile.com/c/lkE0et/esoDJ).

**American DAPs**

In chromosome 6 underrepresented DAP region for the American ancestry one variant (*rs13205210*, chr6:34,831,856) is characterised by a PHRED value of 23.8 and belongs to the protein coding gene UHRF1BP1 which may act as a negative regulator of cell growth.

We found three different regions in chromosome 11 with observed overrepresentation of American ancestry in Caribbean and Puerto Rico (PUR). The first one contains 99 SNPs from 12 genes and has a length of 388kb (chr11:43,908,230-44,296,591). The second one comprises 10 SNPs, with 5 of them with significant allele frequency difference (chr11:46,210,259-46,297,631; 87kb), of which the *rs10437653*  (chr11:46,297,631), has shown association with birth weight in Europeans [[23]](https://paperpile.com/c/lkE0et/WKcka). The third region spans 185kb (chr11:56,000,288-56,184,888) and contains 21 SNPs belonging to 12 genes; the *rs10896290* (chr11:56,128,081) has a PHRED of 23.1 and is part of a gene, OR8J1, and a pseudogene, RPL5P29 (Supplementary Table 5).

**Supplementary figures**

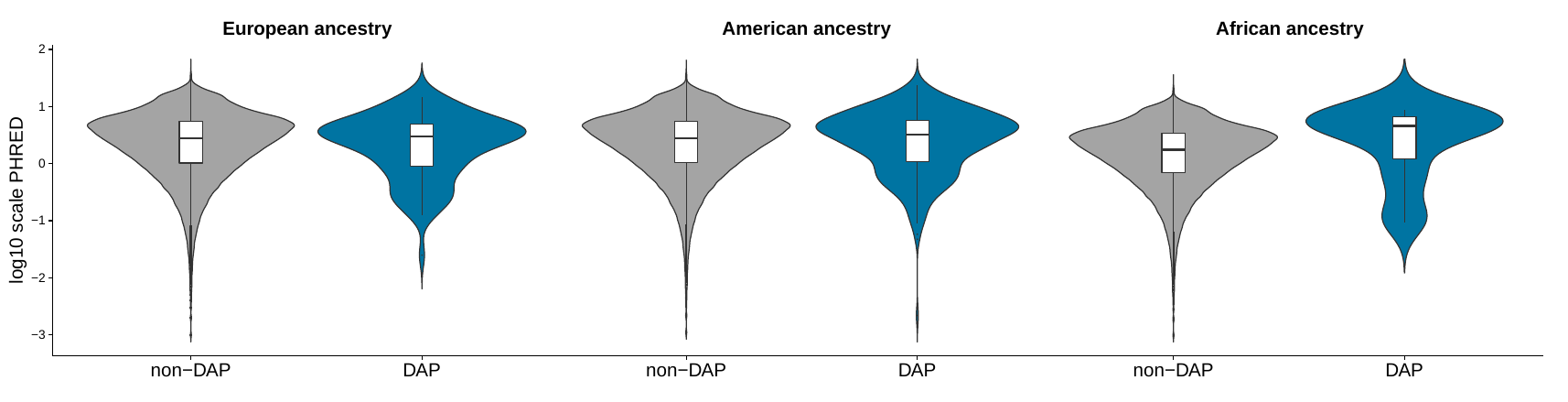

**Supplementary figure 1:** Comparison of the distribution of all the PHRED-scaled C-score values belonging to windows with divergent ancestry profiles (DAP) with the ones from the non-divergent for European (5 DAP and 16,851 non-DAP windows), American (24 DAP and 16,832 non-DAP windows) and African (1 DAP and 16,855 non-DAP windows) ancestries in the 1090Ind dataset. None of the Wilcoxon tests resulted statistically significant (alpha=0.01). The number of analysed windows is reported in Table 6A.

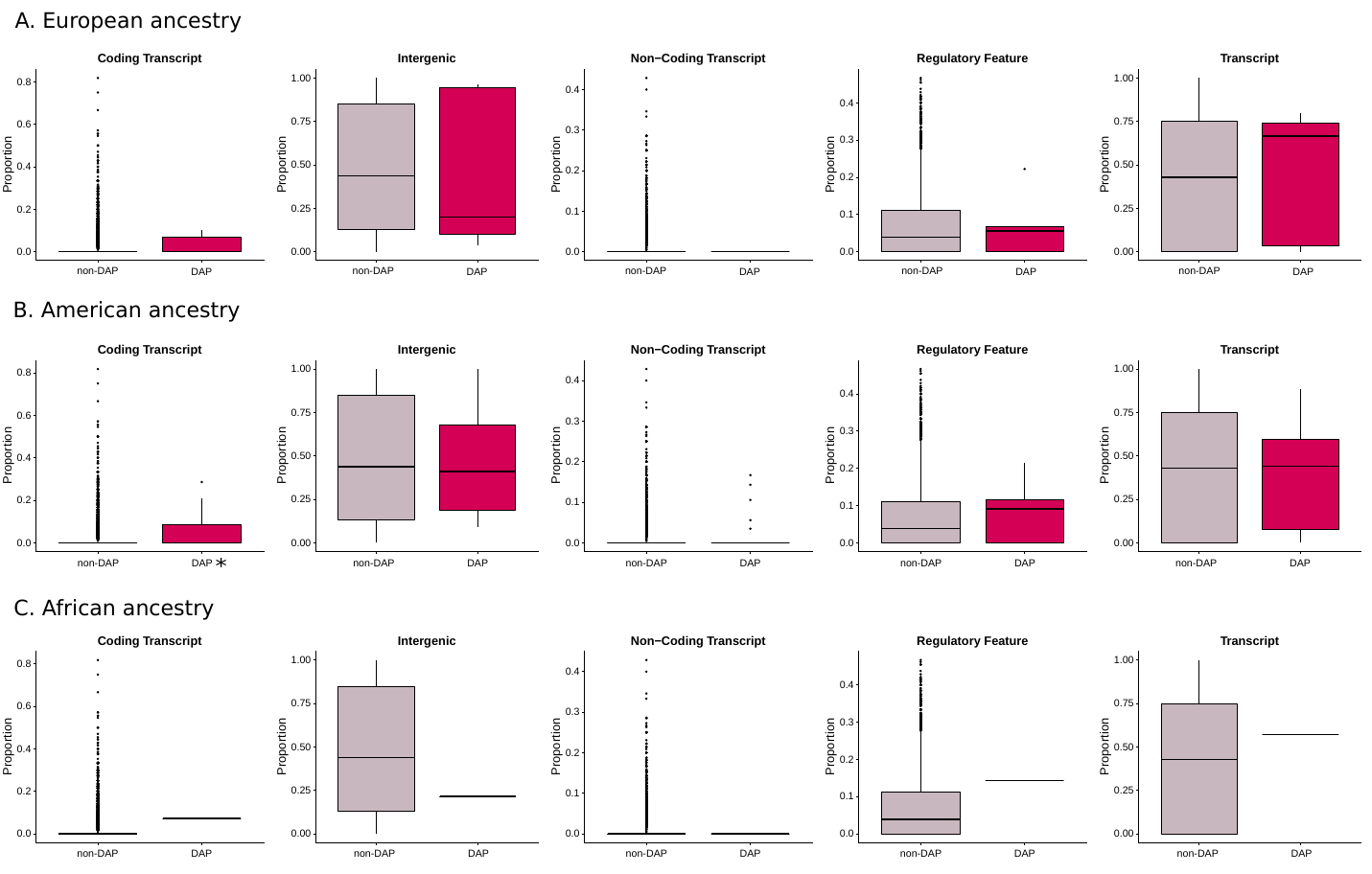

**Supplementary figure 2:** Comparison of the distribution of AnnoTypes (Coding Transcript, Intergenic, Non-coding Transcript, Regulatory Feature, Transcript) belonging to divergent ancestry profiles (DAP) windows with the ones from the non-divergent for European (A), American (B) and African (C) ancestries in the 1090Ind dataset. The asterisk refers to a statistically significant p-value (Wilcoxon test, alpha=0.01). The number of analysed windows is reported in Table 6B.
